# Recombination marks the evolutionary dynamics of a recently endogenized retrovirus

**DOI:** 10.1101/2021.02.24.432774

**Authors:** Lei Yang, Raunaq Malhotra, Rayan Chikhi, Daniel Elleder, Theodora Kaiser, Jesse Rong, Paul Medvedev, Mary Poss

**Author notes:** Corresponding author: Mary Poss. Current address: Department of Hematology and Oncology, University of Virginia, Charlottesville, VA 22903, USA.

## Abstract

All vertebrate genomes have been colonized by retroviruses along their evolutionary trajectory. While endogenous retroviruses (ERVs) can contribute important physiological functions to contemporary hosts, such benefits are attributed to long-term co-evolution of ERV and host because germline infections are rare and expansion is slow, because the host effectively silences them. The genomes of several outbred species including mule deer (*Odocoileus hemionus*) are currently being colonized by ERVs, which provides an opportunity to study ERV dynamics at a time when few are fixed. Because we have locus-specific data on the distribution of cervid endogenous retrovirus (CrERV) in populations of mule deer, in this study we determine the molecular evolutionary processes acting on CrERV at each locus in the context of phylogenetic origin, genome location, and population prevalence. A mule deer genome was de novo assembled from short and long insert mate pair reads and CrERV sequence generated at each locus. CrERV composition and diversity have recently measurably increased by horizontal acquisition of a new retrovirus lineage. This new lineage has further expanded CrERV burden and CrERV genomic diversity by activating and recombining with existing CrERV. Resulting inter-lineage recombinants endogenized and subsequently retrotransposed. CrERV loci are significantly closer to genes than expected if integration were random and gene proximity might explain the recent expansion by retrotransposition of one recombinant CrERV lineage. Thus, in mule deer, retroviral colonization is a dynamic period in the molecular evolution of CrERV that also provides a burst of genomic diversity to the host population.

## Introduction

Retroviruses are unique among viruses in adopting life history strategies that enable them to exist independently as an infectious RNA virus (exogenous retrovirus, XRV) (Coffin 1996) or as an integral component of their host germline (endogenous retrovirus, ERV) (Löwer et al. 1996; Weiss 2006). An ERV is the result of a rare infection of a germ cell by an XRV and is maintained in the population by vertical transmission. Germline colonization has been a successful strategy for retroviruses as they comprise up to 10% of most contemporary vertebrate genomes (Stoye 2012). Over the evolutionary history of the species, ERV composition increases by acquisition of new germ line XRV infections, and through retrotransposition or reinfection of existing ERVs (Boeke and Stoye 1997; Belshaw et al. 2004; Belshaw, Katzourakis, et al. 2005; Johnson 2015), which results in clusters of related ERVs. The ERV profile in extant species therefore reflects both the history of retrovirus epizootics and the fate of individual ERVs. Because the acquisition of retroviral DNA in a host genome has the potential to affect host phenotype (Jern and Coffin 2008; Kurth and Bannert 2010; Feschotte and Gilbert 2012), the dynamic interactions among ERVs and host could shape both retrovirus and host biology. However, the evolutionary processes in play near the time of colonization are difficult to discern based on an ERV colonization event that occurred in an ancestral species. A better understanding of both host and virus responses to recent germ line invasion might inform homeostatic changes in ERV-host regulation that are relevant to the pathogenesis of diseases in which ERV involvement has been implicated (Antony et al. 2011; Magiorkinis et al. 2013; Wildschutte et al. 2014; Li et al. 2015; Li, Yang, et al. 2019; Xue et al. 2020). Fortunately, there is now evidence that retrovirus colonization is occurring in contemporary, albeit often non-model, species (Arnaud et al. 2007; Elleder et al. 2012; Roca et al. 2017), allowing for investigation of ERV dynamics near the time of colonization. Our goal in this research is to investigate the evolutionary dynamics of the phylogenetically distinct ERV lineages that have sequentially colonized mule deer over the approximate million-year history of this species using the complete genome sequence of a majority of coding ERVs in the context of a draft assembly of a newly sequenced mule deer genome.

The life history strategy adopted by retroviruses indicates why this virus family has been so successful in colonizing host germline. Retroviral replication requires that the viral RNA genome be converted to DNA and then integrated into the genome of an infected cell (Coffin et al. 1997). As with many RNA viruses, the virus polymerase enzyme, reverse transcriptase (RT), is error prone, which contributes to a high mutation rate and enables rapid host adaptation. In addition, RT moves between the two RNA copies that comprise a retroviral genome (Luo and Taylor 1990); this process can repair small genomic defects and increases evolutionary rates via recombination if the two strands are not identical. Retroviral DNA is called a provirus and is transported to the nucleus where it integrates into host genomic DNA using a viral integrase enzyme. The provirus represents a newly acquired gene that persists for the life of the cell and is passed to daughter cells, which for XRV are often hematopoietic cells. For a retrovirus infecting a germ cell, all cells in an organism will contain the new retroviral DNA if reproduction of the infected host is successful.

The retroviral life cycle also demonstrates how ERVs can affect host biology (Jern and Coffin 2008; Bolinger and Boris-Lawrie 2009). ERVs require host transcription factors and RNA polymerases to bind to the retrovirus promoter, called long terminal repeats (LTRs), to produce viral transcripts and the RNA genome. Thus, the viral LTRs compete with host genes for transcription factors and polymerases (Sofuku and Honda 2018). A retrovirus encodes at a minimum, genes for the capsid, viral enzymes, and an envelope gene needed for cell entry, which is produced by a sub-genomic mRNA. Hence an ERV also utilizes host-splicing machinery and can alter host gene expression pattern if the site of integration is intronic (Isbel and Whitelaw 2012; Kim 2012). While XRVs are expressed from small numbers of somatic cells, ERVs are present in all cells and ERV transcripts and proteins can be expressed in any cell type at any stage of host development. Hosts actively silence the expression of full or partial ERV sequences by epigenetic methods (Yao et al. 2004; Hurst and Magiorkinis 2017) and by genes called viral restriction factors (Lavie et al. 2005; Matsui et al. 2010; Sze et al. 2013; Bruno et al. 2019; Geis and Goff 2020). Because there will be no record of an ERV that causes reproductive failure of the newly colonized host, ERVs in contemporary vertebrates are either effectively controlled by host actions, are nearly neutral in effects on host fitness, or potentially contribute to the overall fitness of the host (Haig 2012; Göke and Ng 2016; Blanco-Melo et al. 2017; Fu et al. 2019).

The coding portion of a new ERV can be eliminated from the genome through non-allelic homologous recombination (NAHR) between the LTRs, which are identical regions that flank the viral coding portion. A single LTR is left at the site of integration as a consequence of the recombination event and serves as a marker of the original retrovirus integration site (Hughes and Coffin 2004). Most ERV integration sites in humans are solo LTRs (Belshaw, Dawson, et al. 2005; Subramanian et al. 2011). Because the efficiency of NAHR is highest between identical sequences (Hoang et al. 2010), conversion of a full-length ERV to a solo LTR likely arises early during ERV residency in the genome before sequence identity of the LTR is lost as mutations accrue (Belshaw et al. 2007). Because mutations are reported to arise in ERVs at the neutral mutation rate of the host (Kijima and Innan 2010), sequence differences between the 5′ and 3′ LTR of an ERV have been used to approximate the date of integration (Johnson and Coffin 1999; Zhuo et al. 2013).

Although in humans most ERV colonization events occurred in ancestral species, acquisition of new retroviral elements is an ongoing (Stocking and Kozak 2008; Anai et al. 2012) or contemporary (Roca et al. 2017) event in several animal species. The consequences of a recent ERV acquisition are important to the host species because it creates an insertionally polymorphic site; the site is occupied in some individuals but not in others. All ERVs are insertionally polymorphic during the trajectory from initial acquisition to fixation or loss in the genome. Indeed, the HERV-K (human endogenous retrovirus type K) family is insertionally polymorphic in humans (Soriano et al. 1987; Turner et al. 2001; Moyes et al. 2007; Wildschutte et al. 2016) and HERV-K prevalence at polymorphic sites differ among global populations (Li, Lin, et al. 2019). Phylogenetic analyses of the ERV population in a genome can inform on the origins of ERV lineages to determine which are actively expanding in the genome and the mutational processes that drive evolution. These data indicate if expansion is related to the site of integration or a feature of the virus, or both and coupled with information of ERV prevalence at insertionally polymorphic sites, can inform ERV effects on host phenotype.

To this end, we explored the evolutionary history of the mule deer (*Odocoileus hemionus*) ERV (Cervid endogenous retrovirus, CrERV) because we have extensive data for prevalence of CrERV loci in northern US mule deer populations (Bao et al. 2014) and preliminary data on CrERV sequence variation and colonization history (Elleder et al. 2012; Kamath et al. 2013). A majority of CrERV loci is insertionally polymorphic in mule deer; 90% of animals shared fewer than 10 of approximately 250 CrERV integrations per genome in one study (Bao et al. 2014). Further, mule deer appear to have experienced several recent retrovirus epizootics with phylogenetically distinct CrERV and, because none of the CrERV loci occupied in mule deer are found in the sister species, white-tailed deer (*Odocoileus virginianus*) (Elleder et al. 2012), all endogenization events have likely occurred since the split of these sister taxa. Based on the phylogeny of several CrERV identified in the mule deer genome, at least four distinct epizootics resulted in germ line colonization (Kamath et al. 2013). A full-length retrovirus representing the youngest of the CrERV lineages was recovered by co-culture on human cells, indicating that some of these CrERV are still capable of infection (Fábryová et al. 2015). In this study, we expand on these preliminary data by sequencing a mule deer genome and conducting phylogenetic analyses on a majority of reconstructed CrERV genomes. Our results demonstrate that expression and recombination of recently acquired CrERV with older CrERV have increased CrERV burden and diversity and consequently have increased contemporary mule deer genome diversity.

## Results

### Establishing a draft mule deer reference genome to study CrERV evolution and integration site preference

We developed a draft assembly of a mule deer genome from animal MT273 in order to determine the sequence at each CrERV locus for phylogenetic analyses and to investigate the effect of CrERV lineage or age on integration site preference. ERV sequences are available in any genome sequencing data because a retrovirus integrates a DNA copy into the host genome. However, there is extensive homology among the most recently integrated ERVs making them difficult to assemble and causing scaffolds to break at the site of an ERV insertion (Chaisson et al. 2015). We assembled scaffolds using a combination of high coverage Illumina short read whole genome sequencing (WGS) and long insert mate pair sequencing. Our *de novo* assembly yielded an ~3.31 Gbp draft genome with an N50 of 156 Kbp (Table S1), which is comparable to the 3.33 Gbp (c-value of 3.41 pg) experimentally-determined genome size of reindeer (*Rangifer tarandus*) (Vinogradov 1998; Gregory 2019).

Approximately half of CrERV loci are located at the ends of scaffolds based on mapping our previously published junction fragment sequences (Bao et al. 2014), which is consistent with the fact that repetitive elements such as ERVs break scaffolds. To determine the sequence of these CrERVs and the genome context in which they are found, we developed a higher order assembly using reference assisted chromosome assembly (RACA) (J. Kim et al. 2013). RACA further scaffolds our *de novo* mule deer assembly into ‘chromosome fragments’ by identifying synteny blocks among the mule deer scaffolds, the reference species genome (cow), and the outgroup genome (human) (Figure 1A). We created a series of RACA assemblies based on scaffold length to make efficient use of all data (Table S1). RACA150K takes all scaffolds greater than 150,000 bp as input and yielded 41 chromosome fragments, 35 of which are greater than 1.5 Mbp; this is consistent with the known mule deer karyotype of 2n=70 (Gallagher et al. 1994). However, RACA150K only incorporates 48% of the total assembled sequences (1.59 Gbp) because of the scaffold size constraint. In contrast, RACA10K uses all scaffolds 10,000 bp or longer and increases the assembly size to 2.37 Gbp (~72% of total assembly) but contains 658 chromosome fragments (Table S1). The majority of scaffolds that cannot be incorporated into a RACA assembly are close to the ends of alignment chains (File S1, section 1a). Most sequences not represented in any assemblies were repeats based on *k-mer* analyses (File S1, section 1a and Figure S1).

**Figure 1.**
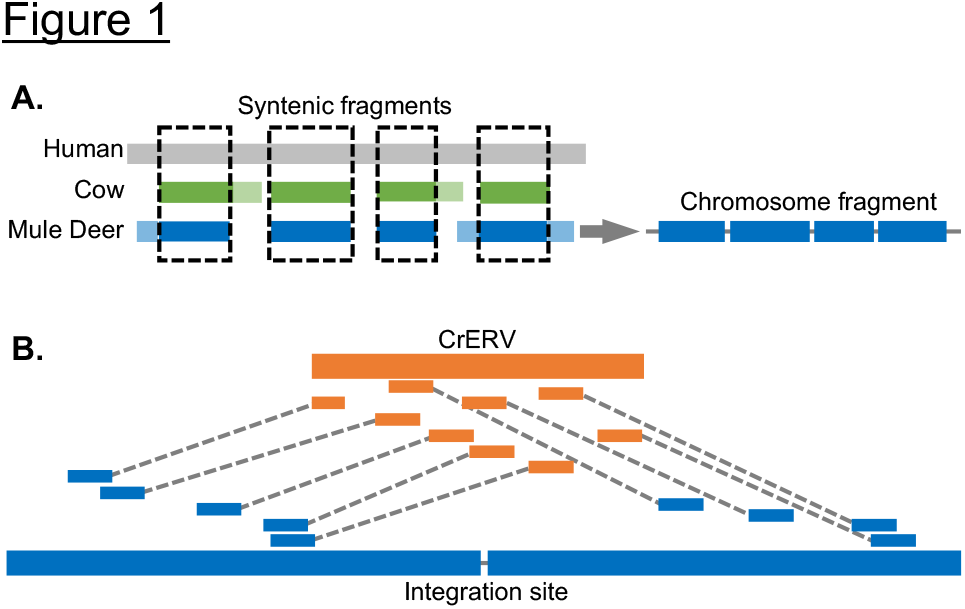
Diagram of CrERV reconstruction and RACA. A. Mule deer chromosome fragment reconstruction using syntenic fragments. Gray, green and blue boxes correspond to aligned human, cow and mule deer scaffold respectively. Lighter shades represent regions that can only be aligned between two species. Dashed boxes highlight syntenic fragments where the region is conserved among all three species, which yield a chromosome fragment that orients mule deer scaffolds. B. Reconstruction of CrERV sequences. CrERV and mule deer scaffolds are shown in bold orange and blue boxes, respectively. Long insert mate pair reads are connected by dotted lines and are colored to indicate whether they derive from the mule deer scaffold or CrERV genome. CrERV genomes were assembled by gathering the broken mate pairs surrounding each CrERV loci as described.

Some scaffolds were excluded from the RACA assemblies, presumably because there is no synteny between cow and human for these sequences. We oriented these scaffolds using the cow-mule deer and sheep-mule deer alignments (RACA+, Table S2). Approximately 124 Mbp of sequence (~4% of total assembly) is in scaffolds larger than 10kb but cannot be placed in RACA10K, nearly all of which can be found on the mule deer-cow alignment chain and the mule deer-sheep (oviAri3) alignment chain (123 Mbp in each chain). Because there is overlap between these alignments, only ~1 Mbp is specific to cow and ~1 Mbp is specific to sheep. Therefore, RACA+ incorporated all but 69 scaffolds that are greater than 10 kbp, which consisted of 1.17 Mbp of sequence (~0.04% of total scaffold size of the assembly) and yields an assembly size of 2.49 Gbp (Table S1).

To enable the investigation of CrERV integration site preference relative to host genes, we annotated the mule deer scaffolds. We used Maker2 (Cantarel et al. 2008; Holt and Yandell 2011) for the annotation, which detects candidate genes based on RNA sequencing data and protein homology to any of the three reference genomes: human, cow and sheep. After four Maker iterations, 21,598 genes with an AED (annotation edit distance) (Cantarel et al. 2008) of less than 0.8 were annotated (Table S3). Approximately 92% of genes are found on RACA150K scaffolds and 95% of genes are represented in RACA10K scaffolds.

### Establishing the location and sequence at CrERV loci

Several lines of evidence suggest that most CrERVs are missing from the assemblies. Only three CrERVs with coding potential were assembled by the *de novo* assembly. The *k-mer* based analysis shows that less than 9.62% of all LTR repeat elements are in the assemblies (Table S4). The CrERV-host junction fragments previously sequenced (Bao et al. 2014) support that CrERV loci are near scaffold ends or in long stretches of ‘N’s. Therefore, we took advantage of the different chromosome fragments generated by RACA10K, RACA150K and RACA+ and the long insert mate pair sequencing data to reconstruct CrERVs at each locus (Figure 1B). We identified 252 CrERV loci in the MT273 genome, which is consistent with our estimates of an average of 240 CrERV loci per mule deer by quantitative PCR (Elleder et al. 2012) and 262 CrERV loci in animal MT273 by junction fragment analysis (Bao et al. 2014). The majority of CrERV loci (206/252) contains CrERVs with some coding capacity and 46 are solo LTRs. Of the 206 CrERVs containing genes, 164 were sufficiently complete to allow phylogenetic analysis on the entire genome or, if a deletion was present, on a subset of viral genes; at 42 loci we were unable to obtain sufficient lengths of high-quality data for further analyses.

### Evolutionary history of CrERV

We previously showed that mule deer genomes have been colonized multiple times since the ancestral split with white tailed deer approximately one million years ago (MYA) (Kamath et al. 2013) because none of the CrERV integration sites are found in white-tailed deer. To better resolve the colonization history, we conducted a coalescent analysis based on an alignment spanning position 1,477-8,633 bp (omitting a portion of *env*) of the CrERV genome (GenBank: JN592050) using 34 reconstructed CrERV sequences with high quality data that had no signature of recombination and that were representative of the phylogeny in a larger data set (Figure 2). The majority of the *env* gene, which has distinct variable and conserved region (Benit et al. 2001), was manually blocked because of alignment difficulties (6,923-7,503 bp by JN592050 coordinates; see Figure 2, right panel for diagram of *env* variable regions and Table S5, column C for *env* structure of each CrERV). This tree shows four well-supported CrERV lineages, each diverged from a common ancestor at several points since the split of mule deer and white-tailed deer. Although *env* sequence is not included in the phylogenetic analysis, CrERV assigned to each of the four identified lineages share the same distinct *env* variable region structure of insertions and deletions, which define the receptor-binding domain of the envelope protein (Figure 2, right panel).

**Figure 2.**
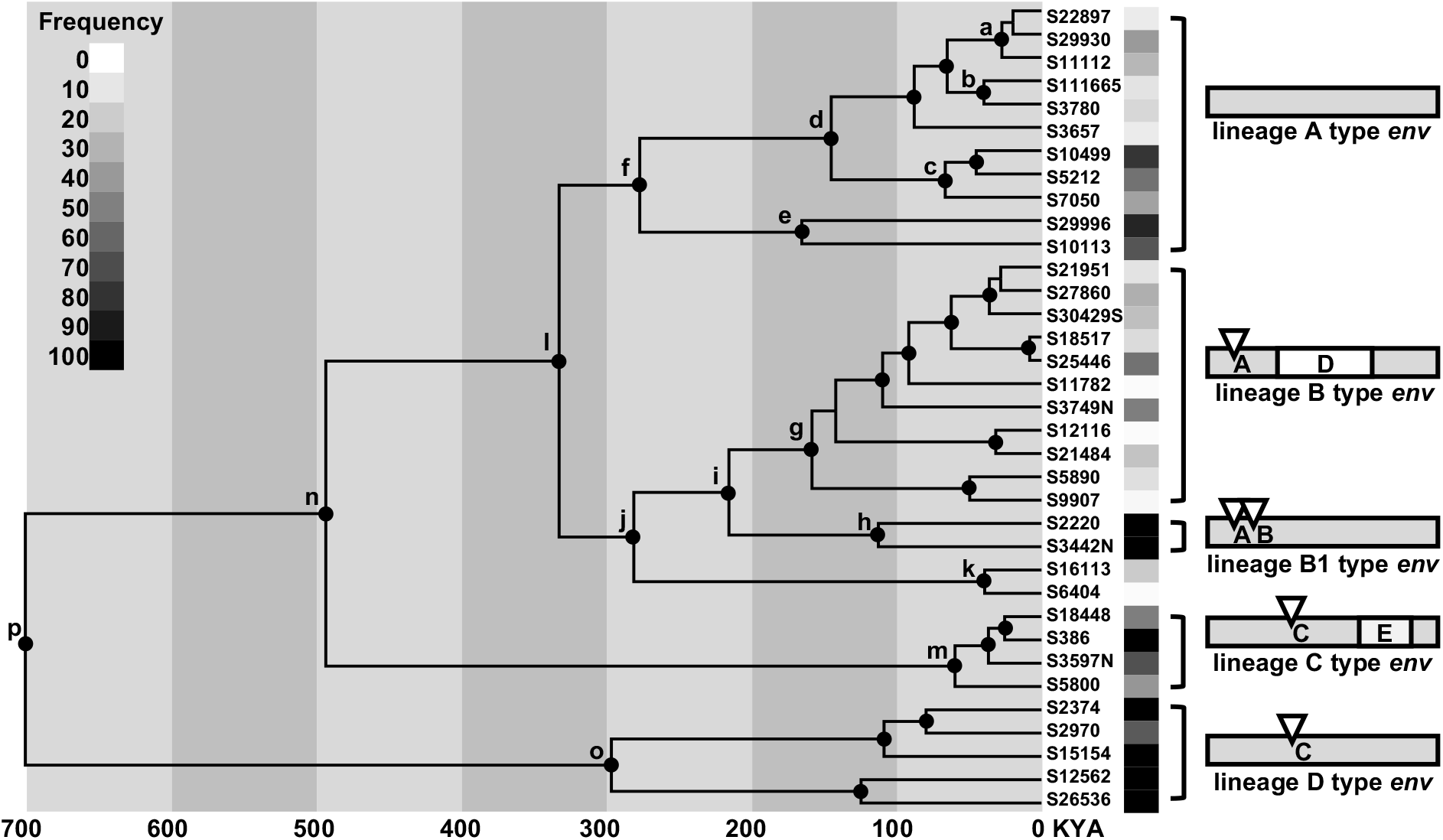
Coalescent phylogeny, *env* structural variation and population frequency of representative full-length non-recombinant CrERVs. Nodes with at least 95% posterior probability support are marked by black dots. The high posterior density for each labeled node is shown in Table S6. Boxes next to CrERV names display the frequency of the CrERVs in the mule deer population with a gray scale (annotated at the top-left corner). Diagrams on the right side depict the lineage-specific structural variations in the CrERV envelope gene. White triangles represent insertions (A, B, C), and white rectangles represent deletions (D and E).

Lineage A CrERVs are the youngest ERV family in mule deer. Our estimates indicate that Lineage A colonization has occurred over the last 300 thousand years to the present (Figure 2; Table S6, node f, 95% high posterior density (HPD) interval 110-470 thousand years ago (KYA)) and is represented by three well-supported CrERV subgroups evolving over this time frame. All have a complete open reading frame (ORF) in *env* and likely represent a recent retrovirus epizootic. An infectious virus recovered by co-culture belongs to this lineage (Elleder et al. 2012). Lineage A represents 30% of all CrERV sampled from MT273 (Table S5). Our age estimates for each subgroup of Lineage A CrERV are consistent with their prevalence in populations of mule deer in the Northern Rocky Mountain ecosystem (Figure 2); (Hunter et al. 2017). For example, S29996 and S10113 are estimated to derive from an older Lineage A CrERV subgroup and occur in our sampled mule deer at higher prevalence than those estimated to have entered the genome more recently (see S22897 and S111665, Figure 2).

Lineage B CrERV shared a common ancestor with Lineage A approximately 300 KYA (node i, Figure 2). Lineage B CrERVs have a short insertion in the 5′ portion of *env* followed by a deletion that removes most of the *env* surface unit (SU) relative to Lineage A *env*. Because our coalescent analysis does not include *env* sequence, these results suggest that two phylogenetically distinct XRV with different envelope proteins were circulating about the same time in mule deer populations. Lineage B CrERV represent 32% of sampled viruses from our sequenced genome (Table S5). Like Lineage A, the prevalence of CrERV from Lineage B among mule deer in the northern Rockies region is low, reflecting their more recent colonization of the mule deer genome. Indeed, six Lineage B CrERVs were identified only in MT273, while only one Lineage A CrERV is found only in MT273 (Table S5), which could be indicative of a recent expansion of some Lineage B CrERV. Of note, there are two related groups of CrERV affiliated with Lineage B (Lineage B1 and B2, Table S5). One shares the short 5′ insertion in *env* but has a full-length *env* with an additional short insertion relative to the *env* of Lineage A CrERV (Lineage B1, Figure 2). CrERV with this *env* configuration represent 9% of coding CrERV in MT273. Because the prevalence of Lineage B1 is high in mule deer, this group could represent the ancestral state for Lineage B CrERVs. The second group has a unique *env* not found in any other CrERV lineages (Lineage B2, Figure 2, node k; S16113 and S6404). We are unable to estimate the prevalence of this unusual *env* containing CrERV in mule deer because the host junction fragments are not represented in our draft mule deer assembly. It is possible that these viruses represent a cross-species infection and it would be interesting to determine if representatives of Lineage B2 are found in the genomes of other species that occupied the ecosystem in the past.

Our coalescent estimates indicate that Lineage C CrERV emerged about 500 KYA (Table S6). Several members of this lineage are found in all mule deer sampled (Figure 2; Table S5), consistent with a longer residence in the genome. There is a 59 bp insertion (C) and 362 bp deletion (E) in *env* (Figure 2; Table S5) compared to the full length *env* of Lineage A; none have an intact *env* ORF. Despite evidence that Lineage C is an older CrERV, the approximately 13% of identified CrERV in MT273 belonging to this lineage share a common ancestor ~50 KYA (95% HPD: 16-116 KYA, Table S6). These data are consistent with a recent expansion of a long-term resident CrERV.

The first representatives of the CrERV family still identifiable in mule deer colonized shortly after their split from white-tailed deer, approximately one MYA (Elleder et al. 2012). Lineage D CrERVs comprise 12% of reconstructed CrERV in MT273 and appear to be near fixation. Indeed, all mule deer in a larger survey of over 250 deer had CrERV S26536, which is not found in white-tailed deer (Kamath et al. 2013). This lineage shares an *env* insertion with Lineage C but lacks the deletion, which removes the transmembrane region of *env*.

These data expand our previous findings that over the approximately one million year history of mule deer, the mule deer genome has been colonized at least four times by phylogenetically distinct CrERVs; this likely reflects several retroviral epizootics each characterized by a unique *env* structure. The two lineages responsible for most recent endogenization events comprise 62% of sampled CrERV. In addition, these data capture the evolutionary processes acting on the *env* gene of exogenous retroviruses, which are characterized by gain or loss of variable regions of this important viral protein.

### Recombination among CrERV lineages

Our coalescent estimates (Figure 2) indicate that two phylogenetically distinct CrERV lineages have been expanding in contemporary mule deer genomes over the last 100,000 years. Both lineages have been actively colonizing contemporary mule deer genomes based on divergence estimates, which include zero. While CrERVs represented by Lineage A are capable of infection (Elleder et al. 2012), all Lineage B CrERVs have an identical deletion of the SU portion of *env* and should not be able to spread by reinfecting germ cells. However, the mule deer genome is comprised of approximately equal percentages of Lineage B and Lineage A CrERVs so we considered two modes by which defective Lineage B CrERVs could expand in the genome at a similar rate with Lineage A. Firstly, ERVs that have lost *env* are proposed to preferentially expand by retrotransposition (Gifford et al. 2012) because a functional envelope is not necessary for intracellular replication. Secondly, we consider that Lineage B CrERVs could increase in the genome by infection if the co-circulating Lineage A group provided a functional envelope protein, a process called complementation (MAGER and FREEMAN 1995; Belshaw, Katzourakis, et al. 2005). This latter mechanism requires that a member of each CrERV lineage be transcriptionally active at the same time in the same cell, and that intact proteins from the ‘helper’ genome be used to assemble a particle with a functional envelope for reinfection. If two different CrERV loci are expressed in the same cell, both genomes could be co-packaged in the particle. Because the reverse transcriptase moves between the two RNA genomes as first strand DNA synthesis proceeds, evidence of inter-lineage recombination would support that the molecular components necessary for complementation were in place. We assessed Lineage B CrERV for recombination with Lineage A to determine if coincident expression of the RNA genomes of these two lineages could explain the expansion by infection through complementation of the *env*-less Lineage B CrERV.

There is good support for recombination between Lineage A and B in a region spanning a portion of *pol* to the beginning of the variable region in *env* (4,422-7,076 based on coordinates of JN592050). In this region, several CrERV that we provisionally classified as Lineage B because they carried the prototypical *env* deletion of SU form a monophyletic group that is affiliated with Lineage A CrERV (Figure 3, upper collapsed clade containing orange diamonds). These Lineage B recombinants all share the same recombination breakpoint just 5′ of the characteristic short insertion for these viruses (Figure S2, indicated by “**”; Table S7). In addition, several other CrERVs with Lineage B *env* branch between lineages A and B, indicating that the recombination breakpoints fall within the region assessed (Figure S2). Indeed, the breakpoint in a group of three CrERV is at position 6630 based on coordinates of JN592050, which is near the predicted splice site for *env* at position 6591 (Elleder et al. 2012); this confers an additional 500 bp of the Lineage B *env* on these viruses (Figure S2) resulting in their observed phylogenetic placement. Because recombination between the two retroviral RNA genomes occurs during reverse transcription, our data indicate that both Lineage A and B CrERVs were expressed and assembled in a particle containing a copy of each genome. A functional envelope from Lineage A would therefore have been available for infection. These data support our premise that complementation with a replication competent Lineage A CrERV or CrXRV (cervid exogenous gammaretrovirus, an exogenous version of CrERV) contributes to the 32% prevalence of *env*-deleted Lineage B CrERV in the genome. It is notable that recent retrotransposition of the lineage A-B recombinant CrERVs likely occurred because they are nearly identical and the branches supporting them are short (Figure 3, orange diamonds in the Lineage A type *env* cluster).

**Figure 3.**
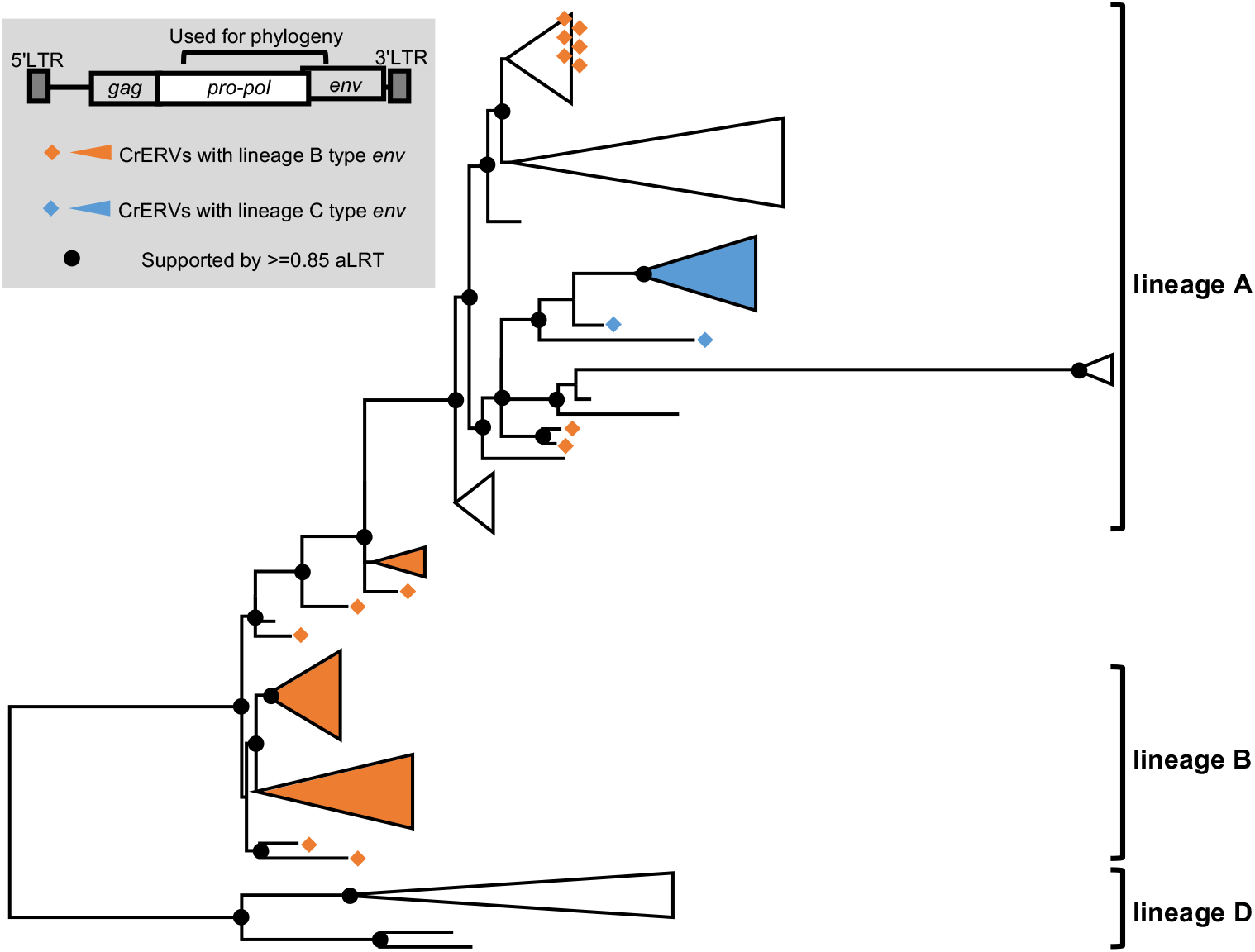
Recombination among CrERVs. Shown is a maximum likelihood phylogeny based on a region spanning a portion of *pol* to *5′env* (JN592050: 4422-7076). Taxa used are a subset of full-length non-recombinant CrERVs representing the four lineages shown in Figure 2 and CrERVs with a recombinant signature containing a Lineage B *env*. Supported nodes (aLRT >= 0.85) are represented by black dots on the backbone of the tree. Lineage designation is assigned to supported branches based on the non-recombinant CrERV. Over this interval, Lineage B CrERVs are found as a sister group to lineage A CrERV but some CrERV containing a prototypical Lineage B *env* are dispersed among Lineage A CrERV. Note that in this interval lineage C CrERVs cluster with Lineage A CrERVs.

There is additional data to support the transcriptional activity of a Lineage B CrERV, which is requisite for recombination with an infectious Lineage A CrERV or for retrotransposition. We identified a non-recombinant Lineage B CrERV (S24870 in Table S5) with extensive G to A changes (184 changes) compared to other members of this monophyletic group. These data are indicative of a cytidine deaminase acting on the single stranded DNA produced during reverse transcription (Suspène et al. 2004).

Lineage C CrERV are enigmatic because based on full length sequences lacking a signature of recombination it diverged around 500KYA (Figure 2) but all extant members of this group diverged recently. From Figure 3, it is evident that over the region of *pol* assessed, CrERVs containing the Lineage C *env* cluster with an older Lineage A subgroup. Given that the *env* of Lineage C CrERV shares sequence homology and an insertion with that of the oldest Lineage D, it is likely that Lineage C is in fact the result of recombination between an early member of Lineage A and a relative of a Lineage D CrERV. Many, but not all, Lineage C CrERVs are found at high prevalence in the mule deer population (Figure 2; Table S5), supporting that the initial recombination event occurred early during the Lineage A colonization. Our identification of Lineage C as derived from a non-recombinant CrXRV is therefore incorrect. Instead, Lineage C CrERVs are derived from a CrERV or CrXRV that is not currently represented in mule deer genomes either because it was lost or it never endogenized. Fourteen of the twenty-two CrERV in Lineage C have multiple signatures of recombination predominantly with Lineage A CrERV. The expansion of a subset of Lineage C as a monophyletic group approximately 50 KYA (Figure 2; Table S6) suggests that like some members of Lineage B, CrERVs generated by recombination with Lineage A have recently retrotransposed.

### Genomic distribution of CrERV lineages

Of the 164 CrERV that we reconstructed from MT273, only 12 can be detected in all mule deer that we have sampled (Kamath et al. 2013; Bao et al. 2014) (Table S5). This means that the majority of CrERV loci in mule deer are insertionally polymorphic; not all animals will have a CrERV occupying a given locus. ERVs can impact genome function in multiple ways but the best documented is by altering host gene regulation, which occurs if the integration site is near a host gene (Rebollo et al. 2012). Thus, we investigated the spatial distribution of CrERV loci relative to host genes to determine the potential of either fixed or polymorphic CrERV to impact gene expression, which could affect host phenotype.

The actual distance between genes is likely to be unreliable in our assembly because most high copy number repeats are missing in the mule deer assembly (Figure S1, Table S4, section 1a of File S1). To investigate potential problems determining the spatial distribution of CrERV insertions imposed by using a draft assembly, we simulated the distribution of retrovirus insertions (File S1, section 2l) in mule deer (scaffold N50=156 Kbp) and the genomes of cow (Btau7, scaffold N50=2.60 Mbp) and human (hg19, scaffold N50=46.4 Mbp). The mean distance between insertion and the closest gene for all simulation replicates (Figure S3) is significantly higher in the cow and human (Mann-Whitney U test *p* < 2.2×10^−16^ for any pair of comparison among the three species). Therefore, we determined if the number of CrERV loci observed to be within 20Kbp of a gene differed from that expected if the distribution was random. There are significantly more observed insertions that fall within 20 Kbp of the translation start site of a gene than occur randomly (Figure 4A). In contrast, intronic CrERV insertions are significantly less than expected based on our simulations (Figure 4B). Among Lineage A CrERVs, only a single sub-lineage (CrERVs that are associated to node ‘a’ in Figure 2) are found in closer proximity to genes (bold font in Column G of Table S5) than expected if integrations are random (Fisher’s exact test *p* = 0.002891). We also investigated whether any of the recombinant CrERV with a signature of recent expansion was integrated within 20 Kbp of a gene. Two of the three recombinant clusters (Figure 3) contain members that are close to a gene (Table S5, bold font in column G). In particular, Lineage A/B recombinant CrERV S10 is 494 bp from the start of a gene. Remarkably, four Lineage C CrERVs with the typical *env* sequence are within 20 Kbp of a gene (Table S5, bold font in column G). Our data indicate that integration site preference overall favors proximity to genes but that this is not reflected in all lineages. In particular, the history of Lineage C CrERV suggests they could have acquired a different integration site preference through recombination that facilitated recent genome expansion.

**Figure 4.**
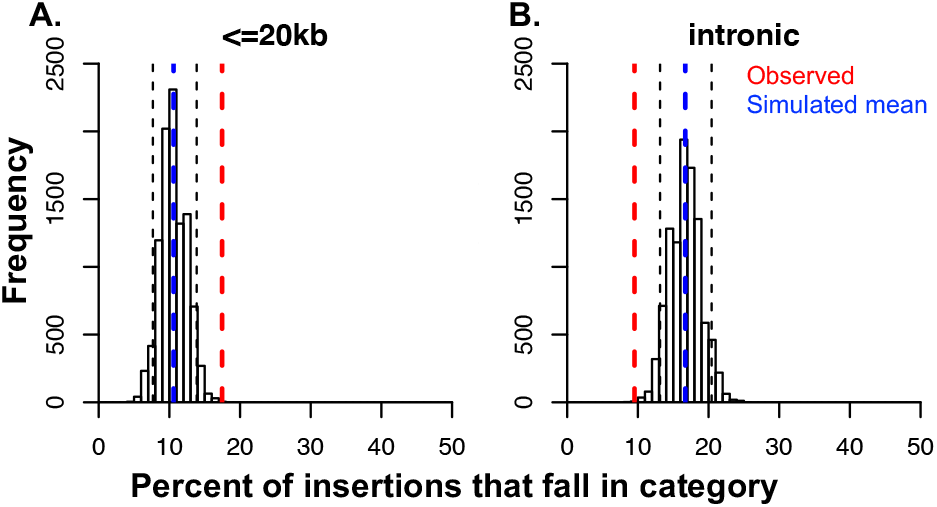
CrERV insertions are enriched within 20 kbp of genes and depleted in introns. We simulated the expected number of CrERV insertions by randomly placing them on the *de novo* assembled MT273 genome. The proportion of insertions expected within 20kb of a gene from the 10,000 replicates is shown in Panel A. The proportion of intronic insertions is in panel B. The distribution of insertions within 20kb of a gene or an intron from the simulation is shown as a histogram. Blue dashed lines indicate the mean of the simulated data. Red dashed lines indicate the observed data in MT273. Black dashed lines indicate the 5th and 95th percentile of the simulated data, which are used to call significant differences.

## Discussion

The wealth of data on human ERVs (HERVs) provides the contemporary status of events that initiated early in hominid evolution. Potential impacts of an ERV near the time of colonization on a host population is thought to be minimal because infection of host germ line by an XRV is a rare event and ERVs that affect host fitness are quickly lost. Potentially deleterious ERVs that are not lost due to reproductive failure can be removed by recombination leaving a solo LTR at the integration site or can suffer degradation presumably because there is no benefit to retain function at these loci; most HERVs are represented by these two states. In addition, humans and other vertebrate hosts have invested extensive genomic resources (Feschotte and Gilbert 2012; Stoye 2012; Zheng et al. 2012) to control the expression of ERVs that are maintained. The dynamics between host and ERV are described as an evolutionary arms race (Daugherty and Malik 2012; Duggal and Emerman 2012). This narrative may underrepresent any contributions of ERVs to fitness as they were establishing in a newly colonized host population. Because there are now several species identified to be at different points along the evolutionary scale initiated by the horizontal acquisition of retroviral DNA it is possible to investigate dynamics of ERV that are not yet fixed in a contemporary species. Considering the numerous mechanisms by which newly integrated retroviral DNA affect host biology, such as by introducing new hotspots for recombination (Campbell et al. 2014), altering host gene regulation (Maksakova et al. 2006; Cohen et al. 2009; Rebollo et al. 2012), and providing retroviral transcripts and proteins for host exaptation (Bénit et al. 1997; Finnerty et al. 2002; Lu et al. 2014; Kawasaki and Nishigaki 2018), colonizing ERVs could make a substantive contribution to species’ evolution. Our research on the evolutionary dynamics of mule deer CrERV demonstrates that genomic CrERV content and diversity increased significantly during a recent retroviral epizootic due to acquisition of new XRV and from endogenization and retrotransposition of recombinants generated between recent and older CrERVs. These data suggest that CrERV provide a pulse of genetic diversity, which could impact this species’ evolutionary trajectory.

Our analyses of CrERV dynamics in mule deer are based on the sequence of a majority of coding CrERVs in MT273. Of the 252 CrERV loci identified in the MT273 assembly, we were able to reconstruct CrERV sequences from long insert mate pair and Sanger sequencing to use for phylogenetic analysis at 164 sites; 46 sites were solo LTR and 42 were occupied by CrERV retaining some coding capacity. We complimented phylogenetic analyses with our previous data on the frequency of each CrERV locus identified in MT273 in a population of mule deer in the northern Rocky Mountain ecosystem (Bao et al. 2014; Hunter et al. 2017). In addition, we incorporated information on the variable structure of the retroviral envelope gene, *env*, which is characteristic of retrovirus lineages but was excluded from phylogenetic analyses. The variable regions of retroviral *env* result from balancing its role in receptor-mediated, cell specific infection while evading host adaptive immune response (Stamatatos et al. 2009; Murin et al. 2019). Despite excluding most of *env* from our phylogenic analysis because of alignment problems, each of the lineages we identified has a similar distinct *env* structure, as is well known for infectious retroviruses. By integrating population frequency, coalescent estimation, and the unique structural features of *env* we provide an integrated approach to explore the evolutionary dynamics of an endogenizing ERV.

The most recent CrERV epizootic recorded by germline infection was coincident with the last glacial period, which ended about 12 KYA. The retroviruses that endogenized during this epizootic belong to Lineage A, have open reading frames for all genes and have been recovered by co-culture as infectious viruses (Fábryová et al. 2015). There are several sub-lineages within Lineage A, which likely reflect the evolutionary history of CrXRV contributing to germline infections over this time period. Lineage A retroviruses constitute approximately one third of all retroviral integrations in the genome. Only four of the fifty Lineage A CrERV that we were able to reconstruct did not have a full length *env*. An important implication of this result is that over the most recent approximately 100,000 years of the evolution of this species, the mule deer genome acquired up to half a megabase of new DNA, which introduced new regulatory elements with promoter and enhancer capability, new splice sites, and sites for genome rearrangements. Thus, there is a potential to impact host fitness through altered host gene regulation even if host control mechanisms suppress retroviral gene expression. None of the Lineage A CrERV is fixed in mule deer populations (Table S5, column F) so any effect of CrERV on the host will not be experienced equally in all animals. However, none of the Lineage A CrERV is found only in M273 indicating that the burst of new CrERV DNA acquired during the most recent epizootic has not caused reproductive failure among mule deer. These data demonstrate that in mule deer, a substantial accrual of retroviral DNA in the genome can occur over short time spans in an epizootic and could impose differential fitness in the newly colonized population.

Lineage A CrERV has an open reading frame for *env* but Lineages B-D do not. Lineage B CrERVs are intriguing in this regard because they also constitute approximately a third of the CrERV in the genome. Yet all have identical deletions of the extracellular portion of *env*, which should render them incapable of genome expansion by reinfection. ERV that have deleted *env* are reportedly better able to expand by retrotransposition (Gifford et al. 2012), which could account for the prevalence of Lineage B. However, because we have evidence for recent expansion of Lineage A and B recombinants, we considered an alternative explanation; that *env*-deficient Lineage B CrERV was complemented with an intact Lineage A CrERV envelope glycoprotein allowing for germline infection. Complementation is not uncommon between XRV and ERV (Hanafusa 1965; Evans et al. 2009), is well established for murine Intracisternal A-type Particle (IAP) (Dewannieux et al. 2004) and has been reported for ERV expansion in canids (Halo et al. 2019). Complementation requires that two different retroviruses are co-expressed in the same cell (Ali et al. 2016). During viral assembly functional genes supplied by either virus are incorporated into the virus particle and either or both retroviral genomes can be packaged. Because the retroviral polymerase uses both strands of RNA during reverse transcription to yield proviral DNA, a recombinant can arise if the two co-packaged RNA strands are not identical. We investigated the possibility of complementation by searching for Lineage A-B recombinants. Our data show that Linage A and B recombination has occurred several times. A group of CrERV that encode a Lineage B *env* cluster with Lineage A CrERV in a phylogeny based on a partial genome alignment (JN592050: 4422-7076bp). The recombinant breakpoint within this monophyletic group is identical, suggesting that the inter-lineage recombinant subsequently expanded by retrotransposition. Notably, two of the CrERV in this recombinant cluster were only found in M273, indicating that retrotransposition was a recent event. There are other clusters of CrERV with Lineage B *env* affiliated with Lineage A CrERV that have different breakpoints in this partial phylogeny. Recombination between an XRV and ERV is also a well-documented property of retroviruses (Kozak 2014; Bamunusinghe et al. 2016; Löber et al. 2018). However, the recombinant retroviruses that result are typically identified because they are XRV and often associated with disease or a host switch. Our data indicate that multiple recombination events between Lineage A and B CrERV have been recorded in germline; this in itself is remarkable given that endogenization is a rare event. Thus both the burden of CrERV integrations and the sequence diversity of CrERV in the mule deer genome increase concomitant with a retrovirus epizootic by CrERV inter-lineage recombination.

Recombination is a dominant feature of CrERV dynamics and is also displayed in the evolutionary history of Lineage C CrERV. Our phylogenetic analysis places the ancestor of Lineage C CrERV at 500 KYA and indeed, Lineage C and Lineage D, which is estimated to be the first CrERV to colonize mule deer after splitting from white-tailed deer (Elleder et al. 2012; Kamath et al. 2013), share many features in *env* that are distinct from those of Lineage A and B. Consistent with a long-term residency in the genome, many Lineage C CrERV are found in most or all mule deer surveyed. A recent expansion of a CrERV that has been quiescent in the genome since endogenizing could explain the estimated 50 KYA time to most recent common ancestor of extant members of this lineage. Although this scenario is consistent with the paradigm that a single XRV colonized the genome and recently expanded by retrotransposition, our analysis shows that all Lineage C CrERV are recombinants of a Lineage A CrERV and a CrERV not recorded in or lost from contemporary mule deer genomes. Hence the resulting monophyletic lineage does not arise from retrotransposition of an ancient colonizing XRV. Rather, as is the case with Lineage B CrERV, recombination between an older CrERV and either a Lineage A CrXRV or CrERV occurred, infected germline, and recently expanded by retrotransposition. It is noteworthy that all retrotransposition events detectable in our data involve recombinant CrERV. Further, recombination often leads to duplications and deletions in the retroviral genome, therefore some of the deletions we document in Lineages B-D are not a consequence of slow degradation in the genome but rather are due to reverse transcription and as was recently reported for Koala retrovirus (Löber et al. 2018).

These data highlight that expansion of CrERV diversity and genomic burden has occurred in the recent evolutionary history of mule deer by new acquisitions, complementation, and pulses of retrotransposition of inter-lineage recombinants. Indeed, several of the recombinant Lineage C CrERVs that have expanded by retrotransposition are within 20kbp of a gene raising the question as to whether there is a fitness effect at these loci that is in balance with continued expression of the retrovirus. It is remarkable that so many of the events marking the dynamics of retrovirus endogenization are preserved in contemporary mule deer genomes. Given that germline infection is a rare event, it is likely that the dynamics we describe here also resulted in infection of somatic cells. It is worthwhile to consider the potential for ERVs in other species, in particular in humans where several HERVs are expressed, to generate novel antigens through recombination or disruptive somatic integrations that could contribute to disease states.

## Materials and Methods

### Sequencing

Whole genome sequencing (WGS) was performed for a male mule deer, MT273, at ~30x depth using the library of ~260 bp insert size, ~10x using the library of ~1,400-5,000 bp insert size and ~30x using the library of ~6,600 bp insert size. 3′ CrERV-host junction fragment sequencing was performed as described by Bao *et al.* (Bao et al. 2014). 5′ CrERV-host junction fragment sequencing was performed on the Roche 454 platform, with a target size of ~500bp containing up to 380 bp of CrERV LTR.

### Assembly and mapping

The draft assembly of MT273 was generated using SOAPdenovo2 (Luo et al. 2012) (File S1, section 2a). WGS data were then mapped back to the assembly using the default setting of bwa mem (Li and Durbin 2009) for further use in RACA and CrERV reconstruction. RNA-seq data was mapped to the WGS scaffolds using the default setting of tophat (Trapnell et al. 2009; D. Kim et al. 2013). 3′ junction fragments were clustered as described in Bao *et al.* (Bao et al. 2014). 3′ junction fragment clusters and 5′ junction fragment reads were mapped to the WGS assembly using the default setting of blat (Kent 2002). A perl script was used to filter for the clusters or reads whose host side of the fragment maps to the host at its full length and high identity. 5′ junction fragments were then clustered using the default setting of bedtools merge.

### RACA

Synteny based scaffolding using RACA was performed based on the genome alignment between the mule deer WGS assembly, a reference genome (cow, bosTau7 or Btau7), and an outgroup genome (hg19). Genome alignments were performed with lastz (Harris 2007) under the setting of ‘--notransition--step=20’, and then processed using the UCSC axtChain and chainNet tools. The mule deer-cow-human phylogeny was derived from Bininda-Emonds *et al.* (Bininda-Emonds et al. 2007) using the ‘ape’ package of R.

### CrERV sequence reconstruction

CrERV locations and sequences were retrieved based on junction fragment and long insert mate pair WGS data. The long insert mate pair WGS reads were mapped to the reference CrERV (GenBank: JN592050) using bwa mem. Mates of reads that mapped to the reference CrERV were extracted and then mapped to the WGS assembly using bwa mem. Mates mapped to the WGS assembly were then clustered using the ‘cluster’ function of bedtools. Anchoring mate pair clusters on both sides of the insertion site were complemented by junction fragments to localize CrERVs. Based on the RACA data, CrERVs that sit between scaffolds were also retrieved in this manner. CrERV reads were then assigned to their corresponding cluster and were assembled using SeqMan (DNASTAR). Sanger sequencing was performed to complement key regions used in CrERV evolutionary analyses. All reconstructed CrERV sequences used in the phylogenetic analyses are included in File S2 in fasta format.

### CrERV evolution analyses

CrERV sequences of interest were initially aligned using the default setting of muscle (Edgar 2004), manually trimmed for the region of interest, and then re-aligned using the default setting of Prank (Löytynoja and Goldman 2005). Lineage-specific regions are manually curated to form lineage-specific blocks. Models for phylogeny were selected by AICc (Akaike Information Criterion with correction) using jModelTest (Posada 2008). Coalescent analysis and associated phylogeny (Figure 2) was generated using BEAST2 (Bouckaert et al. 2014). In the coalescent analysis, we used GTR substitution matrix, four Gamma categories, estimated among-site variation, Calibrated Yule tree prior with ucldMean ucldStddev from exponential distribution, relaxed lognormal molecular clock, shared common ancestor of all CrERVs 0.47-1 MYA as a prior (Elleder et al. 2012; Kamath et al. 2013). Maximum likelihood phylogeny in Figure 3 was generated using PhyML (Guindon and Gascuel 2003) using the models selected by AICc and the setting of ‘-o tlr -s BEST’ according to the selected model.

### CrERV spatial distribution

We simulated 274 insertions per genome to approximate the average number of CrERVs in a mule deer (Bao et al. 2014). The simulation was performed 10,000 times on three genomes: the mule deer WGS scaffolds, cow (Btau7) and human (hg19). Distance between simulated insertions and the closest start of the coding sequence of a gene was calculated using the ‘closest’ function of bedtools, and the simulated insertions that overlap with a gene were marked with the ‘intersect’ function of bedtools. Number of simulated simulations that are within 20 Kbp or intronic to a gene was counted for each of the 10,000 replicates. Counts were then normalized by the total number of insertions and plotted using the ‘hist’ function of R.

## Supplementary methods

Methods with extended details are available in File S1.

## Acknowledgements

This work was supported in part by United States Geological Survey 06HQAG0131. The authors would like to thank David Chen for contributions to genome analysis.

## Availability of Data and Materials

The raw sequencing data was deposited in NCBI BioProject PRJNA705069. Other data generated are included in supplementary file and figures.

## Author’s contributions

LY, RM, RC, TK, JR, PM, and MP conducted analyses; LY, RM, DE, MP interpreted data; LY and MP wrote the manuscript.

