## Supplementary material for "Recombination marks the evolutionary dynamics of a recently endogenized retrovirus": Figure S1

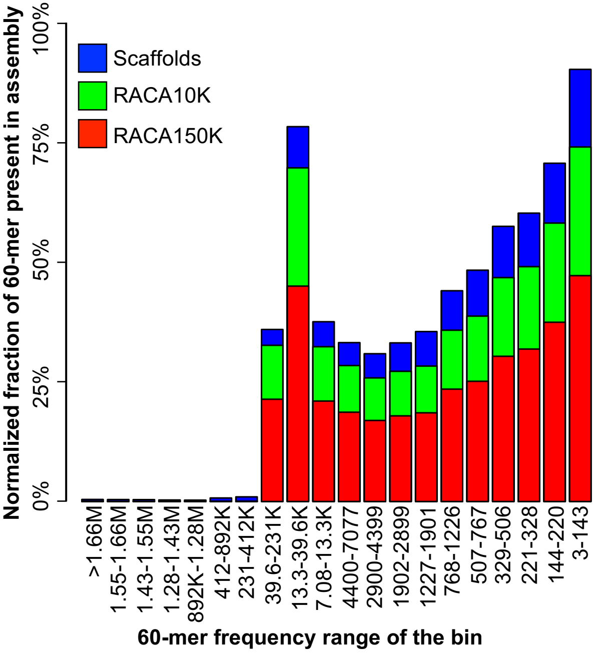


**Figure S1. *K-mer* representation of missing data in assemblies.** *60-mers* were generated from the raw paired-end WGS, then ranked and classified into 20 bins containing equal number of *60-mers*. Sorted by the *60-mer* frequency range, bars represent genome repeats from high copy number (left) to low copy number (right). Color bars show percent of all *60-mers* present in the scaffolds/contigs (blue), RACA10K (green), and RACA150K (red) assemblies. *K-mer* counts were based on the total number of *k-mers* (*k-mer* of frequency *n* were counted *n* times). *K-mers* in the raw sequencing data were normalized based on sequencing depth, genome ploidy, read length and *k-mer* length, so that the *k-mer* fractions reflect the proportion of *k-mers* that are present in each assembly compared to the raw sequencing data. All bars start from 0% instead of being stacked. Numbers beneath each bar indicates the range of frequency of *60-mers* in the raw paired-end genome sequencing data in that bin. M: million, K: thousand.
