## Supplementary material for "Recombination marks the evolutionary dynamics of a recently endogenized retrovirus": Figure S2

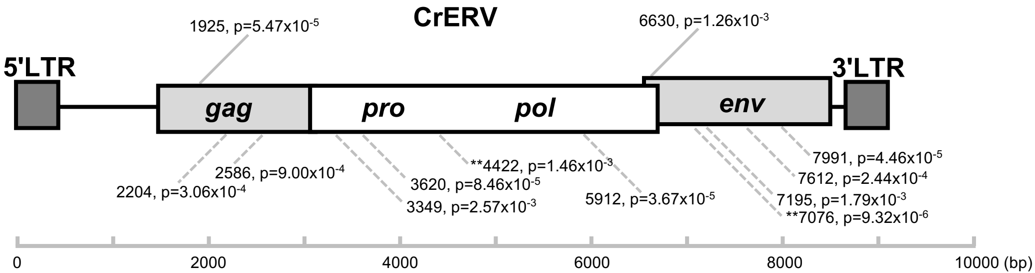


**Figure S2. Diagram of CrERV recombination breakpoints.** Gray lines point at the key recombination breakpoints on the CrERV. Text box connected to the gray lines indicate the coordinate and adjusted p-value of the breakpoint. Solid gray lines indicate breakpoints of recombinant lineages; dashed gray lines indicate additional breakpoints detected by testing the alignment of reference non-recombinant and candidate recombinant CrERVs. All coordinates are relative to GenBank entry JN592050. Double star (**) indicates breakpoints used in the Lineage B recombinant analysis.
