## Supplementary material for "Recombination marks the evolutionary dynamics of a recently endogenized retrovirus": Figure S3

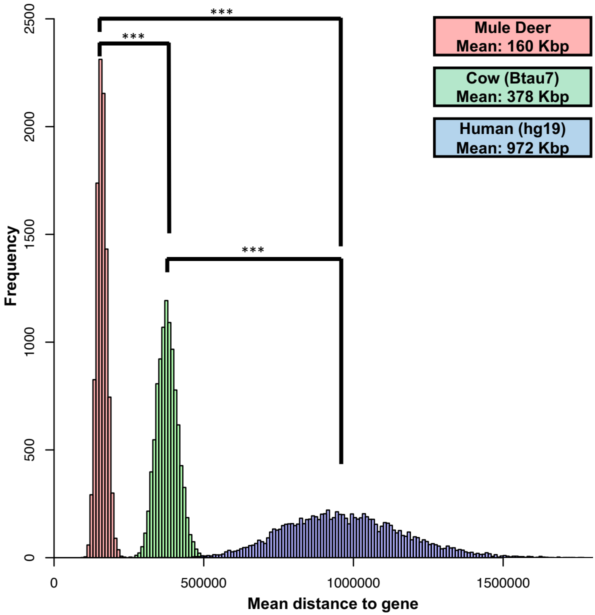


**Figure S3. Distribution of simulated mean distance to gene per replicate in mule deer, cow and human genome.** Distribution of mule deer, cow and human are colored in red, green and blue respectively. Mann-Whitney U test p-values in all three comparisons are less than $2.2\times10^{-16}$.
