## Supplementary material for "Recombination marks the evolutionary dynamics of a recently endogenized retrovirus": File S1

**1. Supplementary analyses**

a. Analysis of non-RACA scaffolds

The mule deer-cow alignment chain still leaves gaps on the cow chromosomes. Ranked by gap size and counted by gap length, the top 5% of RACA-cow alignment gaps are found around the ends of RACA chromosome fragments and repeat-rich regions or assembly gaps of the cow genome. We investigated the repeat nature of unaligned scaffolds using *k-mer* analysis of the raw paired-end WGS, the scaffolds, and each RACA assembly. Table S4 shows the proportion of 60-mers with frequency greater than 50 that are present in each repeat class, which represents sequences with at least 4 copies in the genome given the sequencing depth and *k-mer* length [1]. Taking both total number and fraction missing in consideration, satellite repeat (< 0.15% of raw WGS) and unannotated repeat (< 2.38% of raw WGS) are the most significant contributor of missing repeats in the assemblies. By comparing scaffolds with RACAs, LINE (< 23.35% of raw WGS) and LTR (< 9.62% of raw WGS) elements also contributed considerably to the missing repeat sequences in the RACAs. To evaluate all missing 60-mers, we ranked the 60-mers from the raw paired-end sequencing data by their frequency and categorized them in to 20 bins containing equal total frequency of 60-mers (Figure S1). The frequency of *60-mers* were normalized [1] based on sequencing depth of 23.56x, diploid genome, *k-mer* length 60 and read length of 100 to reflect the proportion of *k-mers* in each bin that are present compared to the raw paired-end sequencing data. The *60-mers* with frequency greater than 231K (or greater than 48K copies in the genome) account for less than 1% of total *60-mers* in their *k-mer* bin, indicating that these high copy number repeats are not assembled. The remaining 13 bins of Figure S1 shows that repeats of lower copy numbers are present in the assemblies at various degrees. The rightmost bin of Figure S1 shows that unique sequences and low copy number repeats (1-30 copies) are mostly present in the assembly (>90%). Furthermore, the lower green and red bars than the blue ones in each bin of Figure S1 shows that RACA10K is less complete than the *de novo* assembly and RACA150K is less complete than RACA10K.

**2. Supplementary methods**

a. Whole genome sequencing, assembly and mapping

Genomic DNA of the lymph node of a male mule deer hunted in Montana (MT273) was extracted using Phenol-Chloroform DNA extraction technique and the library was prepared using Illumina TruSeq DNA PCR free kit. Three libraries were prepared and sequenced: the paired-end library with read length of 2 x 100 bp and target insert size of ~260 bp, sequenced at ~24x depth of a typical mammalian genome; the variable insert length library with read length of 2 x 150 bp and target insert size of ~1,400-5,000 bp averaging at ~3,200 bp, sequenced at ~10x depth of a typical mammalian genome; and the long-insert mate pair library with read length of 2 x 150 bp and target insert size of ~6,600 bp, sequenced at ~24x depth of a typical mammalian genome. Sequencing was performed using Illumina HiSeq 2500 platform at the Pennsylvania State University Genomics Core Facility (https://www.huck.psu.edu/content/instrumentation-facilities/genomics-core-facility).

Before the de novo assembly, paired-end reads were adapter-trimmed using nesoni (https://github.com/Victorian-Bioinformatics-Consortium/nesoni), and both mate-pair libraries were adapter-trimmed using nextclip (https://github.com/richardmleggett/nextclip). The draft assembly was generated using the trimmed reads of all three above mentioned sequencing libraries with SOAPdenovo2 [2] under the setting of 'SOAPdenovo-63mer all -K 61-R -a 400 -p 30'. Further details and discussion that led to the chosen assembly strategy are described at https://www.biostars.org/p/96467/. Untrimmed genome sequencing reads of all three libraries were mapped to the scaffolds using the default setting of bwa mem for further use of RACA assemblies and CrERV reconstructions. Trimming was done to mapped reads per the requirement of downstream analyses.

b. RNA sequencing and mapping

The total RNA from the lymph node of another male Montana mule deer (MT257) is sequenced to aid the annotation of the mule deer genome. Total RNA was extracted using Trizol (Life technologies) and the sequencing library was prepared using the Illumina TruSeq kit. The library was sequenced using the Illumina HiSeq 1500 platform targeting ~10 million 2x36 bp read pairs. Sequencing was conducted at the Pennsylvania State University Genomics Core Facility. RNAseq data was duplicate-removed using the default setting of fastuniq [3], adapter-trimmed using the default setting of Trimmomatic [4], and then mapped to the scaffolds using the default setting of Tophat2 [5,6] and assembled to transcripts using the default setting of cufflinks [7].

c. Mule deer genome annotation

We annotated the mule deer genome using Maker [8,9]. A total of four Maker iterations were performed to obtain the final annotation. The first Maker run was based on evidence of curated human, cow and sheep protein data from the uniprot database (Janurary 2015 release) and the RNA-seq data of a mule deer lymph node. Three additional iterative Maker runs were then conducted based on the first run allowing *ab initio* prediction of genes. Annotated genes are assigned gene symbols following their homologs in the reference species. When an annotated gene and a gene symbol is the 'reciprocal best' alignment of each other in terms of their e-value of the blastx step of Maker, the annotated gene is assigned that gene symbol and marked 'uniq' for high confidence assignment. When an annotated gene’s best-matching gene symbol in blastx does not align to the annotated gene as the best match, the annotated gene is assigned the gene symbol and marked as 'best' to indicate low confidence assignment. To minimize a common problem in annotation where genes span multiple scaffolds [10], we merged the genes that share the same gene symbol and sit on adjacent scaffolds of RACA10k.

d. *K-mer* frequency calculation

*K-mer* frequency tables of the raw paired-end sequencing reads, the SOAPdenovo-assembled scaffolds, the RACA10K, RACA150K were generated using DSK (http://minia.genouest.org/dsk/). For the raw sequencing data, *k-mer* frequency cutoff of 50 was used, and *k-mer* frequency cutoff of 2 were used for all the assembled versions of the genome. Reverse complement *k-mer* pairs were merged and indexed for their frequency in each sequencing and assembly for their frequency with a custom perl script. *K-mers* were annotated using RepeatMasker '-e crossmatch -species mammal -gff -xsmall -qq' using the March 2017 version of the RepBase reference [11]. For each repeat family, *k-mer*s that are found both in the raw paired-end sequence file and the assemblies are counted and presented in Table S4. To assess the repetitiveness in the unassembled genome segments, frequency of all *k-mer*s that are present in the paired-end sequencing data and present in the assemblies are classified into equal-sized bins containing the same number of *k-mer*s by their frequency and visualized (Figure S1).

e. Junction fragment sequencing and mapping

The 3’ CrERV-host junction fragments were sequenced and clustered following Bao *et al*. [12] to localize CrERVs in MT273 and calculate the frequency of reconstructed virus in the mule deer population. We also generated a MT273-specific library targeting the junction between the host and the 5’ end of CrERV. The 5’ junction fragment contains the up to 380 bp of the CrERV LTR and prepped to target read lengths of ~500 bp. This library was sequenced using the Roche 454 platform, targeting ~50,000 of reads with average length of ~500bp. Clustered 3’ CrERV-host junction fragments were mapped to the mule deer scaffolds using blat (http://hgdownload.soe.ucsc.edu/admin/exe/). Only clusters mapping at high identity (>=95%) and full length (less than 5 bp unaligned at the ends of the host section) are kept for further usage. Clusters that mapped multiple times are assigned to the location where the mapping returned the lowest e-value. Clusters that map with a single lowest e-value are taken for further analyses. Clusters that mapped to 2-4 locations with equal lowest e-values are evaluated for their syntenic locations on the cow chromosome using the mule deer-cow alignment chains (refer to the RACA section). If all scaffold targets correspond to the same location on the cow chromosome, the cluster is assigned to the longest scaffold. 5’ CrERV-host junction fragment reads were mapped to the host genome first before clustering. Following the steps used in 3’ junction mapping, an additional step is added to remove the reads that spans the env-3’LTR junction of CrERV, only leaving the true host-5’LTR junction reads. 5’ junction fragment reads that mapped to the same genomic location were merged using bedtools merge [13] and given a score of number of merged reads. Processing of the mapped junction fragments were done with custom perl (https://www.perl.org) scripts.

f. RACA

To prepare for the RACA run and analyses associated with genomic distributions, the draft assembly was aligned human (hg19), cow (bosTau7) and sheep (oviAri3) using lastz [14] under the setting of '--notransition --step=20' excluding the Y chromosome and uncertain contigs in the assembly. Alignments were done to each chromosome of the reference genomes and then combined to gain computation speed. Lastz alignments were further processed to generate UCSC chain and net files using the UCSC axtChain and chainNet tool (http://hgdownload.soe.ucsc.edu/admin/exe/) under default setting. Considering the completeness and relationship to mule deer, RACA was ran using cow as the reference and human as outgroup. A mule deer-cow-human phylogeny was extracted from Bininda-Emonds *et al*. [15] with the 'ape' package (https://cran.r-project.org/web/packages/ape/index.html) of R (https://www.r-project.org/). The mean and variance of the insert size of sequencing libraries were calculated using a custom perl script. The mule deer-cow, mule deer-human chain and net files, the mule deer-cow-human phylogeny, the mapped sequencing reads and their insert size distribution were used as the input of RACA. Using the same input, RACA was ran at 10 Kbp and 150 Kbp resolution separately.

RACA chromosome fragments represent the synteny blocks that are evolutionarily conserved. However, the adjacent scaffolds that emerged after the split of the cow and the mule deer lineage cannot be retrieved by RACA. We oriented the scaffolds that are not incorporated into the RACA chromosome fragments. For each longer than 10 kbp mule deer scaffold, the best alignment chain by score to cow and sheep are extracted. The extracted alignment chains are sorted by their corresponding coordinate on the cow and sheep chromosomes. Scaffolds in RACA chromosome fragments are attached to the corresponding chain, and thus also sorted by the cow and sheep chromosomes. The sorted chain files reveals additional scaffold adjacency with lower confidence compared to RACA chromosome fragments. The analyses mentioned in this paragraph were done using custom perl scripts.

g. CrERV identification and reconstruction

CrERVs in the MT273 genome are identified with mate pair and junction fragment sequencing data. Variable (1,400-5,000 bp) and long (6,500 bp) insert mate pair reads were first mapped to the CrERV reference as mentioned above. Then, the mate of any CrERV-mapping read is mapped to the host genome assemblies and clustered using bedtools cluster [13] requiring distance between reads (-d) to be no larger than 1,497 bp and reads in the cluster to be mapped in the same orientation. These clusters are hence named 'anchoring discordant mate pair cluster' (ADMPC). Anchoring reads that fall into the same ADMPC are processed to remove duplicates and sequencing adapters using fastuniq and trimmomatic. Given the nature of the mate pair sequencing library, a CrERV insertion corresponds to a pair of ADMPC containing reads oriented towards the insertion. Therefore, the ADMPCs containing the most anchoring reads and within 30 kbp of each other and oriented towards each other are paired to support a CrERV insertion. The pairing was done both on the scaffolds and RACA10K, so that CrERVs that sit on scaffold ends are included. The pairing of ADMPCs is done with a custom perl script. The '-d' parameter in bedtools were selected to maximize the density of each ADMPC.

A coding CrERV insertion is called for having an ADMPC with at least two reads. Because a CrERV insertion corresponds to a pair of ADMPCs, adjacent ADMPCs oriented towards each other are paired to support a single CrERV. ADMPC pairing was done on both scaffolds and RACA10K, so that the CrERVs between two scaffold ends can be identified. CrERV-host junction fragments within 10 kbp of ADMPCs are also incorporated to further support CrERV insertion. Only the mule deer-specific CrERV are included in the analysis [16] and identified by mapping to a contemporary CrERV (GenBank accession JN592050). Identity was determined by aligning reads in ADMPCs using the default setting of blat. Only loci where the ADMPC reads have higher average identity to the JN592050 are considered a CrERV integration and used for subsequent analysis. The sequence of a coding CrERV was reconstructed using the reads in the corresponding ADMPCs. Sanger sequencing of PCR products was conducted to cover gaps in contigs mapping to the coding regions and to confirm ambiguous sites. The PCR products constituted full length CrERV generated with PCR primers in the flanking host region. PCRs were performed using GoTaq (Promega) or Phusion (Thermo-Fisher) depending on the size of the amplicon. PCR products were either cloned with the TA cloning kit (Life Technologies) or directly sequenced using sequencing primers targeting the region of interest.

Presence of a CrERV solo LTR is characterized by CrERV-host junction fragments or LTR sequence in the de novo assembly without ADMPCs around. We identified the assembled LTR sequences in the scaffold by aligning the In7 and In1 LTR described in Kamath *et al*. [16] using blat.

h. Alignment of CrERV sequences

Alignments used in this work were done in two steps. Firstly, the complete sequence of reconstructed CrERVs sequences were aligned using the default setting in Muscle [17]. Secondly, alignment sections were retrieved with a custom perl script, and the sequence sections were realigned using the default setting Prank [18]. Alignments were manually inspected and curated to put regions containing lineage-specific structural variations into blocks.

i. Recombination analyses

Recombinant CrERVs were identified by interrogating various CrERV alignments. Initial classification of CrERVs were made based on the structural variations in the envelope gene sharing the same insertions or deletions that form their own blocks in their alignment (Table S5). Sub-alignments of the 1,477-8,633 segment of CrERVs (coordinate relative to GenBank JN592050) were made for CrERVs of each type of envelop. Recombination signal were detected using the 'Phi' function of PhiPack [19] using the default parameter. Within each CrERV *env* type, the alignment of any combination of CrERVs between four and the total number of CrERVs of the 1,477-8,633 segment were tested. The combinations without recombination signal (Phi-test p-value < 0.01 indicates recombination) and the maximum amount of CrERVs were used to make maximum likelihood phylogeny using PhyML following the AICc criterion with jModelTest. Phylogenies were then visualized to select the combination of CrERVs that maximizes number of supported clusters on the tree.

The detected non-recombinant CrERVs (Table S6A) were used as the baseline set to find recombination breakpoints. Recombination breakpoints were detected with the 'Profile' function of PhiPack [19] to scan the alignment with a window size of 1 Kbp and offset of 25bp. Multiple tests using different '-w' parameters between 50 and 450 with an offset of 50 were used in the Phi Profile program to accommodate cases where phylogenetic informative sites are far from each other in the alignment. Non-recombinant CrERVs within each envelope group were tested together to find the breakpoints of recombinant lineages (Table S6B). All full-length CrERVs belonging to the same envelope type were tested to find breakpoints of within or between lineage recombinants (Table S6B). P-values were adjusted using the Bonferroni method to accommodate multiple testing, and a cutoff of 0.01 of the adjusted p-value was used to call significant recombination breakpoint. Key breakpoints used to determine segments used in phylogenetic analyses were plotted in Figure S2.

j. Coalescent phylogeny of CrERV

Coalescent phylogeny was generated using BEAST2 [20]. For the phylogeny shown in Figure 2, representative CrERVs were selected from the non-recombinant CrERVs (refer to the "recombinant analyses" section) from each major lineage with the preference of CrERVs with less missing data and maximizes the CrERV diversity within each lineage. Sub-alignment of CrERVs at 1,477-8,633 bp (coordinate relative to JN592050) was used to minimize missing data and maximize the proportion of CrERV protein-coding genes covered. For the coalescent analysis, used general time reversible nucleotide substitution matrix, estimated proportion of invariable sites, and among-site variation with four Gamma categories according to the model test performed by AICc (Akaike Information Criterion with correction) in jModelTest [21]. We used Calibrated Yule tree prior with ucldMean and ucldStddev sampled from exponential distribution. For molecular clock, a relaxed clock lognormal was used. The common ancestor of all CrERVs in the phylogeny was set to be a uniform distribution between 0.47 to 1 MYA [16,22] as a prior. The non-In1-like CrERVs were set to be monophyletic as a prior. The iterative chain in BEAST2 was let run until the explained sum of squares of all parameters reach 200. Then, the final phylogeny tree was generated using the TreeAnnotator program in the BEAST2 package taking mean heights of the tree and 50% burnin. The tree was plotted using FigTree (http://tree.bio.ed.ac.uk/software/figtree/).

k. Maximum likelihood phylogeny of CrERV

Maximum likelihood phylogenies were used in Figure 3, as well as phylogenetic incongruence analysis during recombination detection. These phylogenies were generated using PhyML [23] based on the alignments generated as described above and model selection tests. Model selection were performed on corresponding alignments using AICc criterion in jModelTest [21]. Node support was calculated using alpha likelihood ratio test. We also used '-o tlr -s BEST' in the PhyML command line to control the tree optimization and topology search.

l. CrERV spatial distribution simulation and analyses

To simulate random insertions in a genome, random numbers ranging from 1 to the size of the target genome was generated using the 'rand' function of perl. Scaffolds or chromosomes of the target genome were assumed to be laid out one by one to cover the full length of the genome, so that random numbers can be converted to a coordinate on scaffolds or chromosomes with using perl scripts. Y chromosome and the unknown chromosome contigs were not included for the cow and human simulations. We simulated 274 insertions per genome to approximate the number of CrERV insertions in an average mule deer [12], and replicated 10,000 times in each genome. The simulation was done in mule deer (scaffolds), cow (Btau7) and human (hg19).

To calculate the distance between simulated insertions and the nearest gene, we used the abovementioned mule deer gene annotation generated by Maker, and the September 2016 version of RefSeq annotations for cow (Btau7) and human (hg19). The start of the first exon was used as a surrogate of the location of a gene. We used the 'intersect' function of bedtools to find the insertions that overlapped with genes. Then, the insertions that do not overlap with genes were fed to the 'closest' function of bedtools to find the closest gene to each simulated insertion in the target genome. For mule deer, an additional step is taken to find the closest gene on R10K and R150K. When an insertion falls on a scaffold without annotated genes, the closest distance calculated based on R10K and R150K is used. For the observed distance to gene in MT273, a custom perl script was used to find the point of the junction fragments and ADMPCs of a CrERV that is the closest to the insertion site, and that point is used as the integration site in the distance to gene calculation. Mean distance to gene, 5th and 95th percentile were calculated using the 'quantile' function of R, the Wilcoxon test were performed with the 'wilcox.test' function of R.

To categorize the insertions by their distance to the closest gene, we counted the number of genes that are within 20 Kbp and intronic to a gene, and repeated this process for al 10,000 simulation and the observed data in MT273. Counts were then normalized to percentage by the total number of simulated insertions in each simulation replicate or observed data. Categorization were performed using custom perl scripts. Distribution of counts by percentage were then generated using the 'hist' function of R.
